# Delineating the heterogeneity of senescence-induced-functional alterations in hepatocytes

**DOI:** 10.1101/2023.09.04.556220

**Authors:** Pavitra Kumar, Mohsin Hassan, Frank Tacke, Cornelius Engelmann

## Abstract

**Background and Aim:** Cellular senescence of hepatocytes involves permanent cell cycle arrest, disrupted cellular bioenergetics, resistance to cell death, and the release of pro-inflammatory cytokines. This ’zombie-like’ state perpetuates harmful effects on tissues and holds potential implications for liver disease progression. Remarkably, senescence exhibits heterogeneity, stemming from two crucial factors: the inducing stressor and the cell type. As such, our present study endeavors to characterize stressor-specific changes in senescence phenotype, its related molecular patterns, and cellular bioenergetics in primary mouse hepatocytes (PMH) and hepatocyte-derived liver organoids (HepOrgs).

**Methods:** PMH, isolated by collagenase-perfused mouse liver (C57B6/J; 18-23 weeks), were cultured overnight in William’s E-medium supplemented with 2% FBS, L-glutamine, and hepatocyte growth supplements. HepOrgs were developed by culturing cells in a 3D matrix for two weeks. The senescence was induced by DNA damage (doxorubicin, cisplatin, and etoposide), oxidative stress (H_2_O_2_, and ethanol), and telomere inhibition (BIBR-1532), p53 activation (nutlin-3a), DNA methyl transferase inhibition (5-azacitidine), and metabolism inhibitors (galactosamine and hydroxyurea). SA-β galactosidase activity, immunofluorescence, immunoblotting, and senescence-associated secretory phenotype (SASP), and cellular bioenergetics were used to assess the senescence phenotype.

**Results:** Each senescence inducer triggers a unique combination of senescence markers in hepatocytes. All senescence inducers, except hydroxyurea and ethanol, increased SA-β galactosidase activity, the most commonly used marker for cellular senescence. Among the SASP factors, CCL2 and IL-10 were consistently upregulated, while Plasminogen activator inhibitor-1 exhibited global downregulation across all modes of senescence. Notably, DNA damage response was activated by DNA damage inducers. Cell cycle markers were most significantly reduced by doxorubicin, cisplatin, and galactosamine. Additionally, DNA damage-induced senescence shifted cellular bioenergetics capacity from glycolysis to oxidative phosphorylation.

**Conclusion:** In our study, we demonstrated that each senescence inducer activates a unique combination of senescence markers in PMH. Doxorubicin demonstrated the highest efficacy in inducing senescence, followed by cisplatin and H_2_O_2_, with no impact on apoptosis. Each inducer prompted DNA damage response and mitochondrial dysfunction, independent of MAPK/AKT.

**Graphical abstract:** 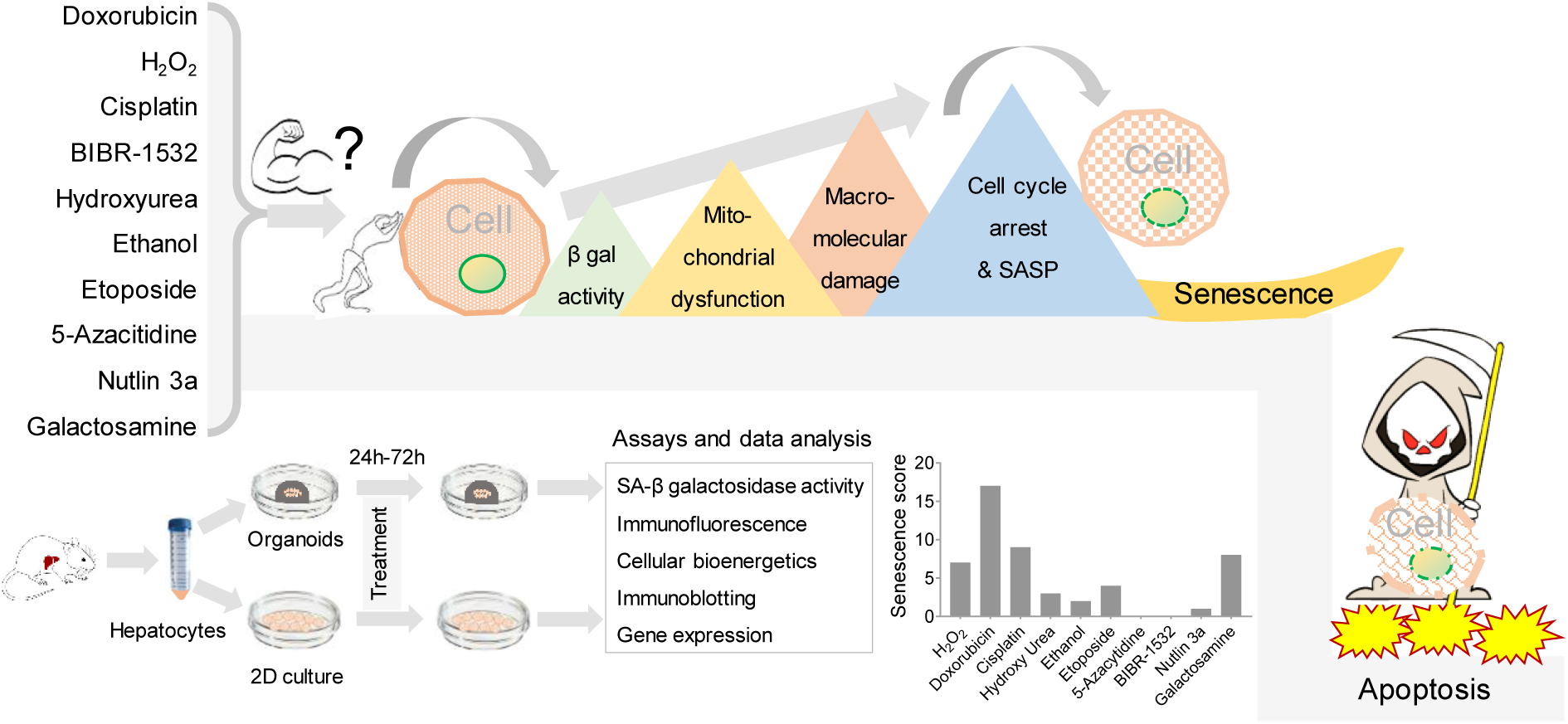

## 1. Introduction

Senescence is an intricate cellular process encompassing permanent cell cycle arrest, metabolic changes, evasion of cell death, and the release of pro-inflammatory cytokines ^1^. The process can be triggered by DNA damage ^2^, oncogene activation ^3^, telomeres shortening ^4^, mitochondrial dysfunction ^5^, and oxidative stress ^6^. The liver, a tissue with robust regenerative capacities, suffers significantly from the accumulation of senescent cells, resulting in a potential reduction of its functionality by over 50% ^7^. Accumulation of senescent hepatocytes may trigger persistent inflammation, fibrosis, and liver cancer ^8 9^ The senescence phenotype is characterized by four main hallmarks: cell cycle withdrawal, macromolecular damage, deregulated metabolism, and the senescence-associated secretory phenotype (SASP) ^10^ all of which hold implications for tissue injury and recovery. Cell cycle withdrawal is assessed by evaluating the expression and activation of cyclins, cyclin-dependent kinases (CDKs), proliferating cell nuclear antigen (PCNA), nuclear membrane integrity, and histones ^11^. Macromolecular damage involves DNA breaks, impaired repair mechanisms, and oxidations of proteins and lipids ^12^. Deregulated metabolism includes dysfunctional mitochondria, impaired lysosomal activity, and glycolysis ^13^. The SASP consists of pro-inflammatory cytokines, chemokines, growth factors, and matrix metalloproteases ^14^.

Despite extensive research, the molecular heterogeneity among senescence cells has posed challenges in characterizing senescence and developing targeted therapies ^15 16^. While senescence-associated β-galactosidase (SA-β-gal) activity is commonly used as a senescence marker, its reliability is debated due to potential activity in non-senescent phases^17^. On the other hand, cell cycles arrest markers such as cyclins, CDKs, lamin B1, and histones show relative consistency across senescence inducers and cell types.

SASP, expressed during the late phase of senescence, is primarily regulated by mitogen-activated protein kinase (MAPK) and NF-κB pathways ^18^. Nevertheless, the composition of SASP varies depending on the mode of senescence induction, stress duration, and cell type. Similarly, while many studies indicate a shift towards glycolysis in senescence ^19 20^, there are also a few reports in contradiction. For instance, DNA damage-induced senescence increased oxidative phosphorylation in human primary dermal fibroblasts, while glycolysis remained unchanged ^21^. Another study by Quijano et al. demonstrated higher rates of both glycolysis and oxidative phosphorylation in oncogene-induced senescent cells ^22^. The substantial heterogeneity among senescent cells can be attributed to two key factors: the type of stressor inducing senescence and the specific cell type undergoing this process. However, senescence heterogeneity and its underlying molecular patterns have not been studied in the hepatocytes. Therefore, it becomes imperative to identify stressor- and cell-type-specific changes in molecular patterns to address the complexity of senescent cell heterogeneity.

Given the diversity among senescent cells, comprehending stressor-specific molecular patterns becomes crucial. In this study, we investigate stressor-specific senescence phenotypes and molecular patterns in primary mouse hepatocytes (PMH). Ten common senescence inducers representing DNA damage, oncogene activation, oxidative stress, telomerase inhibition, and metabolic stress were selected. By studying the effects of these inducers on PMH and hepatocyte-derived organoids, we aim to overcome the limitations of two-dimensional PMH models and gain a deeper understanding of the geometry and replicative aspects of senescence in this context. Through the exploration of PMH’s specific responses to these inducers, we aspire to illuminate the intricate molecular patterns underlying senescence heterogeneity and contribute to the development of targeted therapeutic strategies.

## 2. Results

### 2.1 Differential effects of senescence inducers on hepatocytes’ senescence-associated β-galactosidase activity, p53 and, DNA damage response in primary mouse hepatocytes

To assess the effects of various senescence inducers on SA-β-Gal activity in primary mouse hepatocytes (PMH), we treated the cells with pre-optimized dosages of different senescence inducers (Figure S1 A&B). After optimization of dosage and treatment time, subsequent assays were performed with a 24-hour treatment period and a single dose of each senescence inducer. Apoptosis was assessed using Bax/bcl2 ratio to ensure the non-toxic effect of the drugs. None of the senescence inducers at the optimized dosage and treatment time induced apoptosis, in fact, hydroxy urea, 5-azacitidine, and galactosamine reduced Bax/bcl2 ratio (Figure S2). H_2_O_2_ (oxidative stress), doxorubicin, cisplatin (DNA damage), 5-azacitidine (methyl transferase inhibition, BIBR-1532 (telomere inhibition), and nutlin 3a (p53 activation) increased SA-β-Gal activity significantly. However, hydroxy urea and ethanol did not affect SA-β-Gal activity (Figure 1A).

**Figure 1.**
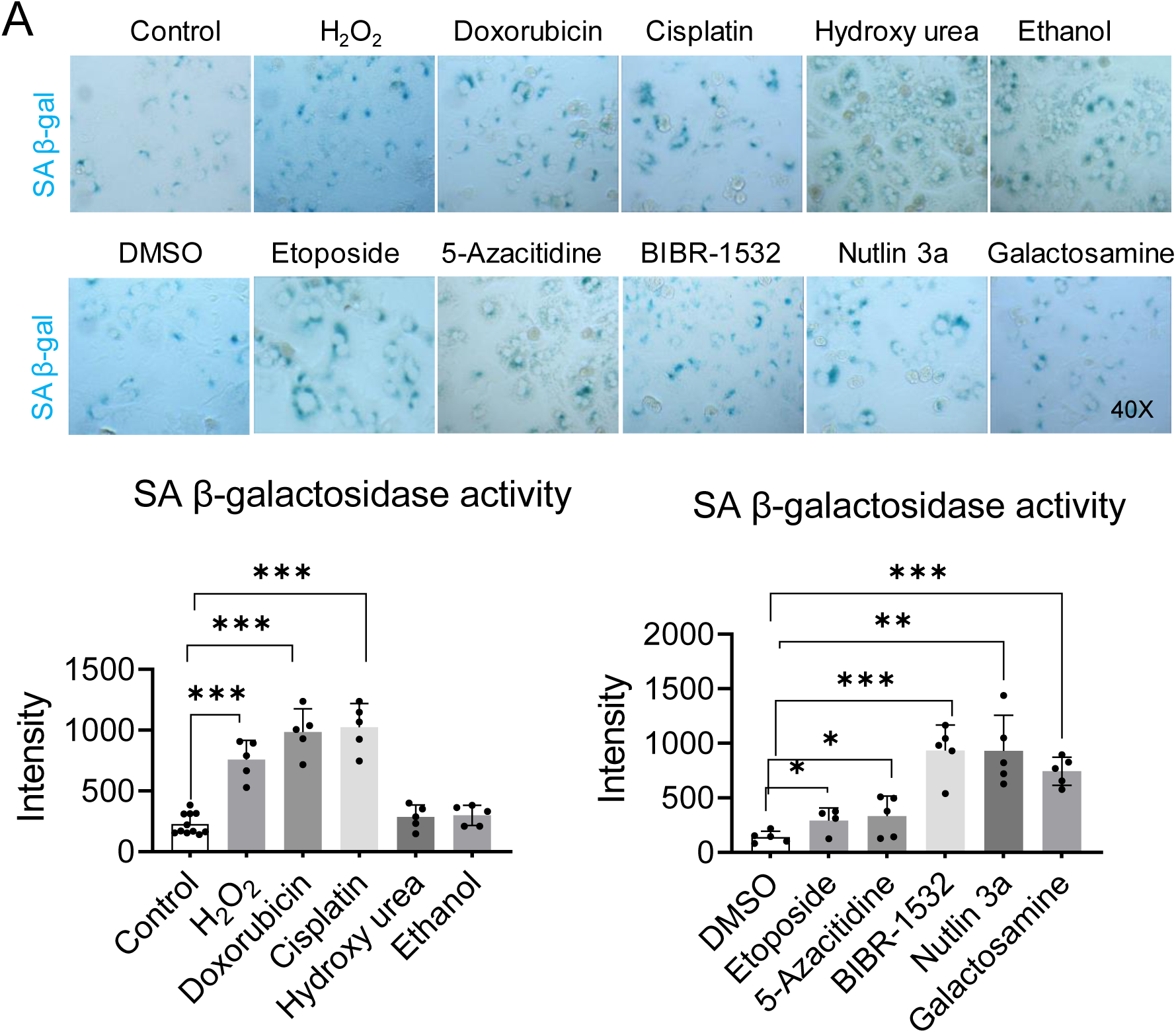
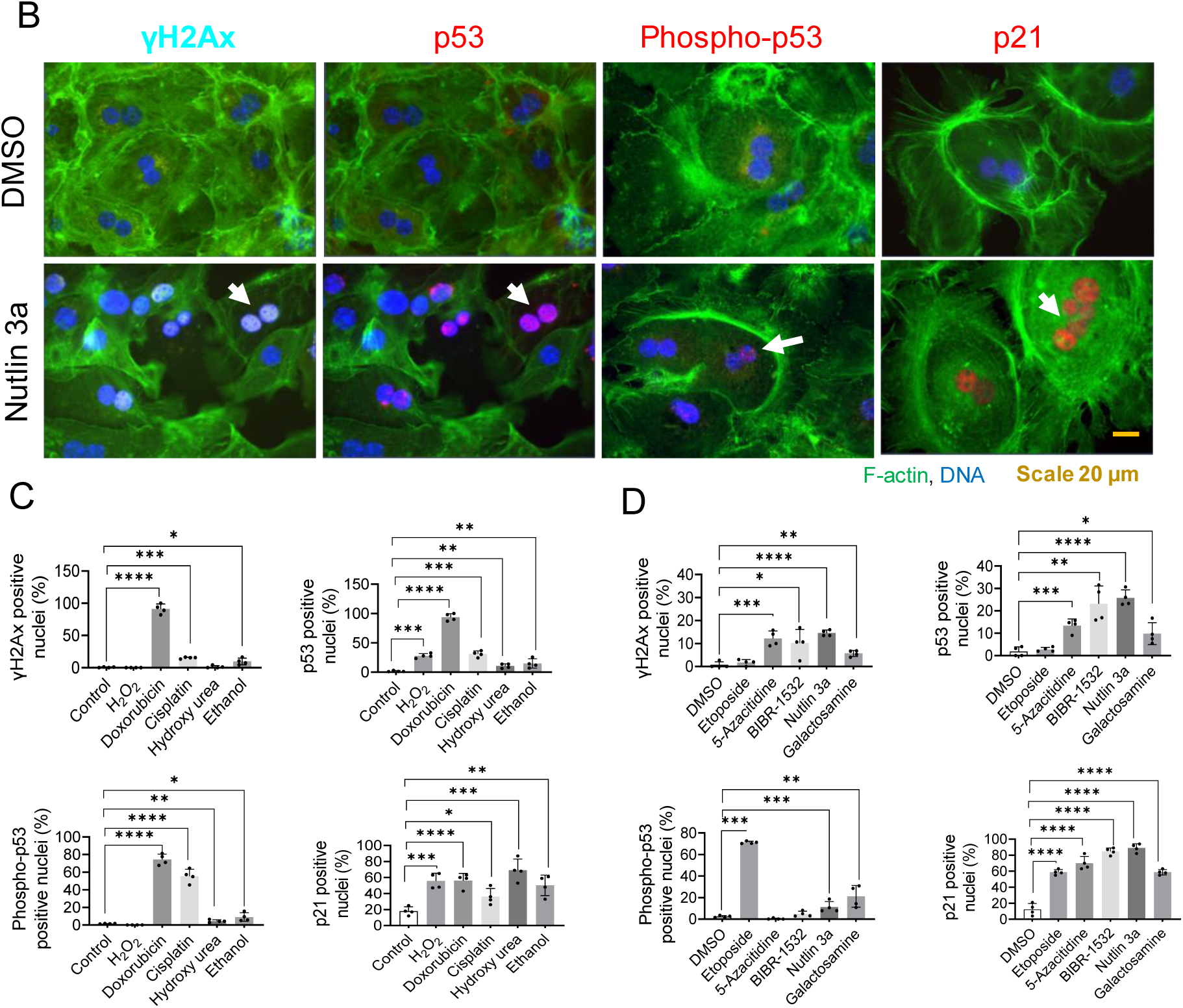
Effects of senescence inducers on hepatocytic senescence-associated β-galactosidase activity, p53 and, DNA damage response in primary mouse hepatocytes. A) Represented images of SA-β-Gal activity after 24 hours of treatment of primary mouse hepatocytes with senescence inducers. The intensity was quantified with Image J. Data were collected from four biological replicates (n=4) and five technical replicates. The mean of technical replicates was used for statistical analysis. Student’s t-test was used to compare the effect of each treatment with the respective control. Error is shown as standard deviation. (*P < 0.05, **P < 0.005, and ***P < 0.001). B) Immunofluorescence images represent senescence-induced DNA damage response markers in primary mouse hepatocytes treated for 24 hours in 2D culture C) Drugs soluble in water and, relative quantification. D) Drugs soluble in DMSO and, relative quantification. Data were collected from four biological replicates (n=4) and five technical replicates. The mean of technical replicates was used for statistical analysis. Student’s t-test was used to compare the effect of each treatment with the respective control. Error is shown as standard deviation. (*P < 0.05, **P < 0.005, ***P < 0.001, and ****P < 0.0001).

Since most of the senescence inducers were able to enhance SA-β-Gal activity, we were interested in analyzing to what extent senescence inducer activates markers of DNA damage response (DDR) and the p53/p21 pathway, which is finally mediating cell cycle arrest. Among the inducers examined, doxorubicin, a potent topoisomerase II inhibitor, proved to be the most effective in inducing phosphorylation of H2AX, a well-established marker of DNA double-strand breaks, and activation of p53. Additionally, under doxorubicin treatment, p21, the downstream target of p53 exhibited significant overexpression. Etoposide, another topoisomerase II inhibitor, did not induce phosphorylation of H2AX; nevertheless, it activated the subsequent p53 pathway. On the other hand, oxidative stress induced by H_2_O_2_ did not trigger the phosphorylation of H2AX and p53, but it did result in the activation of p53, subsequently leading to the activation of p21. Telomerase inhibition by BIBR-1532 and MDM2 inhibition by Nutlin 3a both activated DNA damage and stimulated the p53 pathway. Hydroxyurea, known to impede DNA synthesis through ribonucleotide reductase inhibition, activated p53, thereby upregulating p21. Notably, hydroxyurea did not induce an increase in H2AX phosphorylation, indicating its inability to induce DNA double-stranded breaks. Acute depletion of the uridine pool by Galactosamine increased H2AX phosphorylation and activated p53, followed by an elevation in p21 expression. Similarly, both cisplatin, which cross-links DNA, and 5-azacitidine, which induces DNA hyper-methylation via DNA methyltransferase inhibition, increased H2AX phosphorylation and activated the p53 pathway, leading to heightened p21 expression. Moreover, ethanol exhibited a comparable albeit lesser extent of H2AX phosphorylation and p53 activation, ultimately resulting in increased p21 expression (Figure 1 B-D) (Figure S3 A-B).

### 2.2 Senescence inducers decrease cell cycle progression and proliferation markers

Cell cycle arrest, a prominent hallmark of senescence, was thoroughly investigated to understand the effects of different senescence inducers on cell cycle and proliferation markers. H_2_O_2_ treatment triggered an increase in cdc2 phosphorylation and elevated levels of cyclin A2, indicating an attempt by cells to proliferate. However, this was accompanied by decreased levels of cdc2, lamin B1, Histone 3, and PCNA, suggesting obstacles in the cell cycle and DNA replication processes. In contrast, both doxorubicin and cisplatin induced a strong cell cycle arrest by downregulating cdc2, Cyclin A2, lamin B1, Histone 3, and PCNA, indicating a clear disruption in cell cycle progression. Intriguingly, hydroxyurea and ethanol increased the expression of cdc2, cyclin A2, lamin B1, and PCNA, while simultaneously reducing cdc2 phosphorylation and histone 3 phosphorylation, pointing towards a potential imbalance in cell cycle regulation and DNA replication. Etoposide and 5-azacitidine decreased cdc2 and cyclin A2 levels without affecting cdc2 phosphorylation, lamin B1, histone 3, or PCNA levels. BIBR-1532 exhibited a decrease in cdc2, histone 3, and lamin B1 levels without affecting cdc2 phosphorylation or PCNA levels, suggesting disrupted cell cycle regulation, compromised chromatin organization, and perturbed nuclear envelope maintenance, all of which could contribute to altered cell growth. Notably, both Nutlin 3a and galactosamine led to decreased cdc2 phosphorylation and reduced levels of cdc2, cyclin A2, lamin B1, histone 3, and PCNA, collectively indicating impaired cell cycle processes. These interconnected findings shed light on the intricate relationship between senescence inducers and the regulation of cell cycle progression, emphasizing the complex nature of senescence-associated cellular responses (Figure 2 A-D) (Figure S4).

**Figure 2.**
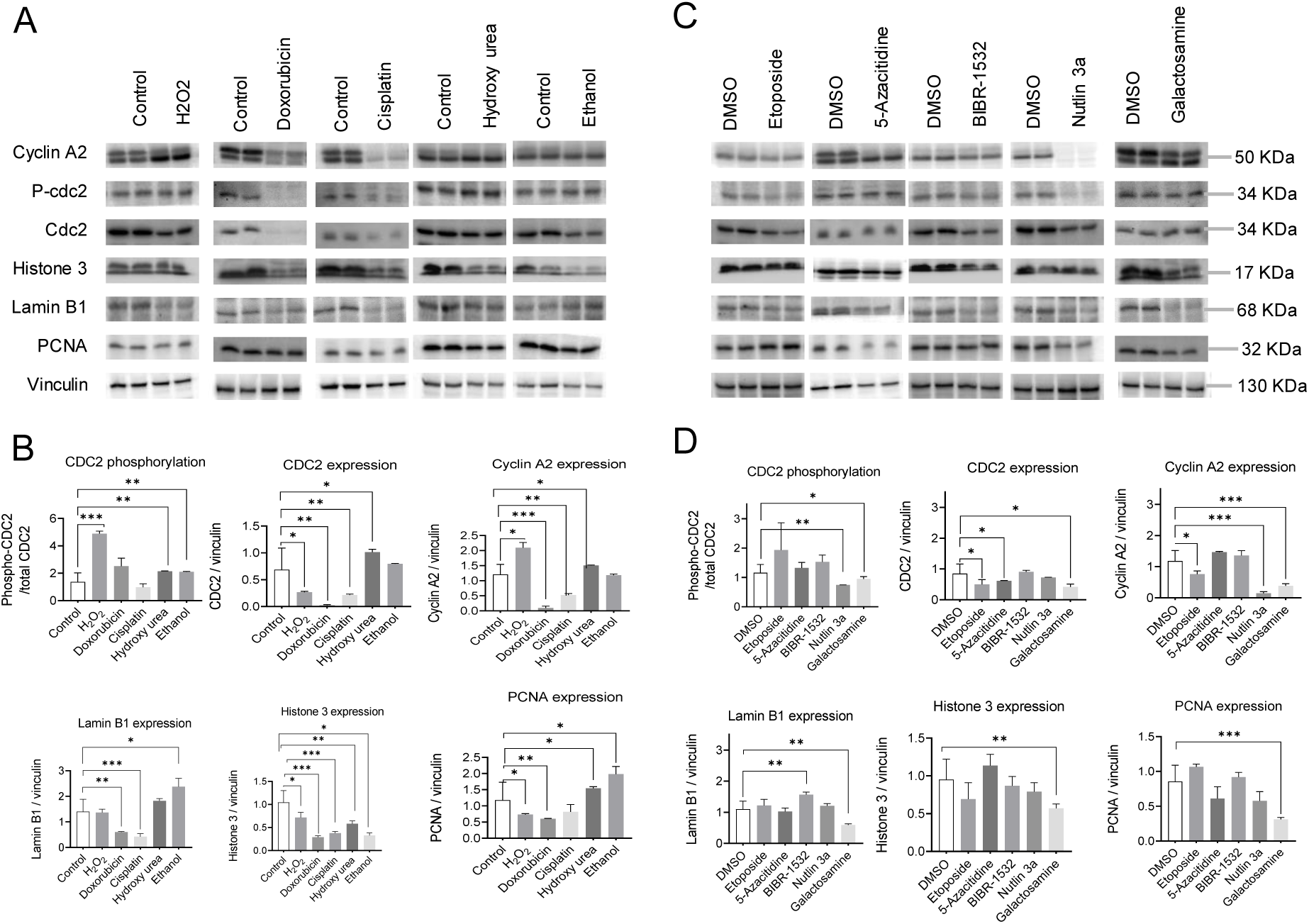
Senescence inducers halt cell cycle progression in primary mouse hepatocytes. Representative Immunoblotting images of the effect of senescence inducers on cell cycle and proliferation markers in primary mouse hepatocytes treated for 24 hours in 2D culture. Vinculin served as a loading control. A) Drugs soluble in water and, B) relative quantification. C) Drugs soluble in DMSO and, D) relative quantification. Data were collected from four biological replicates (n=3). The band intensity was quantified using Image J. Student’s t-test was used to compare the effect of each treatment with the respective control. Error is shown as standard deviation. (*P < 0.05, **P < 0.005, and ***P < 0.001).

### 2.3 Senescence inducers increase the expression of SASP candidates

SASP is a hallmark of senescence leading to immune activation and immune cell recruitment. In order to decipher how different types of senescence inducers may modulate cytokine secretion in hepatocytes, we examined the expression of cytokines and chemokines including C-C Motif Chemokine Ligand 2 (CCL2), C-X-C motif ligand 2 (CXCL2), interleukin 6 (IL6), interleukin 10 (IL10), interleukin 1β (IL1β), plasminogen activator inhibitor-1 (PAI-1), tissue inhibitor of metalloproteinase (TIMP), tumor necrosis factor alpha (TNFα), transforming growth factor beta 1 (TGF1β), and matrix metallopeptidase 13 (MMP13). Interestingly, although doxorubicin and etoposide both target the same enzyme, topoisomerase II, to induce DNA damage, their SASP profiles differed significantly. Both doxorubicin and etoposide upregulated CCL2, TIMP, and MMP13, while IL6, IL10, IL1β, and TNFα were upregulated by doxorubicin but downregulated by etoposide. Doxorubicin alone upregulated CXCL2 and downregulated TGF1β. At the kinase level, doxorubicin activated mitogen-activated protein kinase (MAPK) and mammalian target of rapamycin (mTOR) but deactivated protein kinase B (AKT), whereas etoposide activated AKT and mTOR while deactivating MAPK. Similarly, the DNA cross-linker cisplatin upregulated CXCL2, IL6, IL10, IL1β, and MMP13, while downregulating CCL2, PAI-1, TIMP, and TGF1β by activating MAPK and deactivating AKT. DNA hypermethylation induced by 5-azacitidine resulted in the upregulation of IL6, IL10, TIMP, and the downregulation of CXCL2, IL1β, TNFα, and MMP13 through the deactivation of AKT, MAPK, and mTOR. Furthermore, we investigated the effects of a telomerase inhibitor, BIBR-1532, on SASP expression. Telomerase inhibition upregulated CCL2, IL10, and TIMP while downregulating CXCL2, IL6, PAI-1, and MMP13 by deactivating AKT and MAPK.

Moreover, oxidative stress induced by H_2_O_2_ or ethanol resulted in the upregulation of CCL2, IL6, IL10, TIMP, TGF1β, and MMP13. CXCL2 and TNFα were upregulated specifically by H_2_O_2_, while PAI-1 was downregulated by both H_2_O_2_ and ethanol. IL1β and TNFα, however, were downregulated exclusively by ethanol treatment. At the kinase level, H_2_O_2_ upregulated AKT expression but decreased its phosphorylation, and downregulated mTOR expression, whereas ethanol deactivated AKT, MAPK, and mTOR. Furthermore, interference with the nucleotide pool by galactosamine (which depletes the uridine pool) or hydroxyurea also affected the expression of SASP. Galactosamine upregulated CCL2, CXCL2, IL6, and IL10 while downregulating PAI-1, TIMP, TNFα, and TGF1β through the deactivation of MAPK and mTOR. Hydroxyurea, which depletes deoxyribonucleotides by inhibiting ribonucleoside diphosphate reductase, upregulated CCL2, IL6, IL10, TIMP, and MMP13, and downregulated CXCL2, IL1β, PAI-1, TNFα, and TGF1β by deactivating AKT, MAPK, and mTOR.

Finally, we investigated the activation of p53, a key event in senescence, using Nutlin 3a, an inhibitor of the E3 ubiquitin ligase MDM2. Nutlin 3a upregulated CCL2, IL10, and TIMP, while downregulating IL6, TNFα, TGF1β, and MMP13 by activating AKT and mTOR and deactivating MAPK. These interconnected findings provide valuable insights into the complex regulation of SASP and its modulation by different senescence inducers, highlighting the multifaceted nature of the senescent phenotype (Figure 3 A-B).

**Figure 3.**
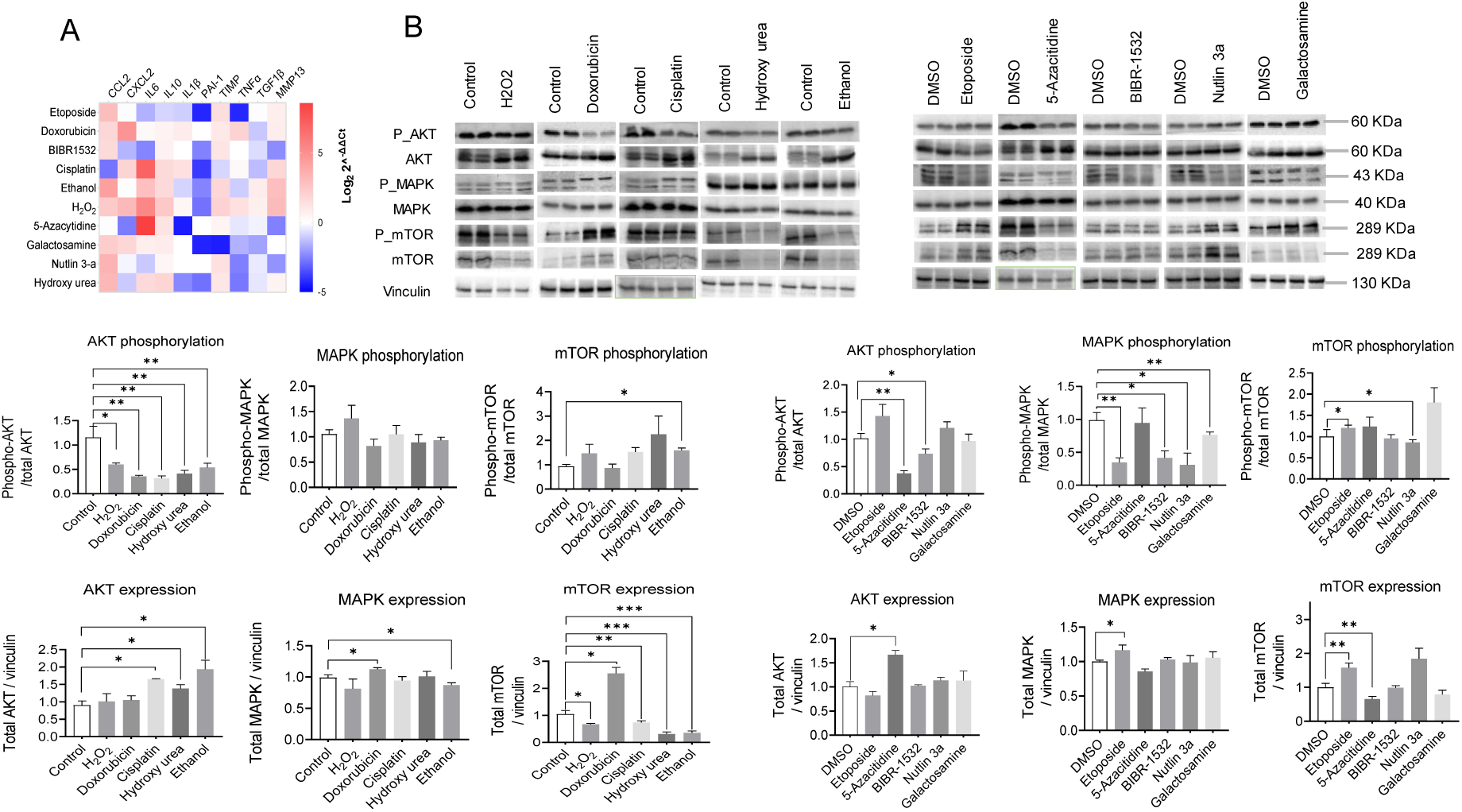
Different senescence inducers in primary mouse hepatocytes dynamically regulate senescence-associated-secretory phenotype (SASP) A) Heat map representing the gene level expression of cytokines, chemokines, and matrix metalloproteases selected among the SASP candidates. GAPDH served as a reference gene and the data show the fold change over the control or DMSO group, as per the vehicle used for the respective drug. The data represent the mean of three biological and two technical replicates. B) Representative Immunoblotting images of the protein kinases regulating SASP expression. Vinculin served as a loading control and the signal was normalized to vinculin probed for each membrane. The band intensity was quantified using Image J. Student’s t-test was used to compare the effect of each treatment with the respective control. Error is shown as standard deviation. (*P < 0.05, **P < 0.005, and ***P < 0.001).

### 2.4 Senescence inducers alter cellular bioenergetics

Cellular metabolic changes are the hallmark of several liver diseases and may contribute to the disease development, notably of non-alcoholic fatty liver disease. Therefore, we sought to investigate the impact of various senescence inducers on cellular bioenergetics. To capture the dynamic changes, we measured cellular bioenergetics at multiple time points ranging from 12 to 72 hours. Comparing the results to the control group, we observed that ethanol initially led to a reduction in both basal and maximum glycolytic and oxidative respiratory capacity. However, intriguingly, at the 48-hour mark, we noted a slight increase in the overall respiratory capacity, which subsequently declined after 72 hours of treatment. Conversely, the introduction of H_2_O_2_ resulted in a profound decrease in the overall respiratory capacity of the cells. On the other hand, the inducers cisplatin, BIBR-1532, and 5-azacytidine initially exhibited an augmentation of oxidative phosphorylation capacity within the first 24 hours. Nevertheless, beyond the 48-hour threshold, the glycolytic capacity became elevated instead. Treatment with Nutlin 3a, conversely, led to a reduction in the overall respiratory capacity of the cells. Interestingly, hydroxyurea administration induced a significant increase in glycolytic capacity while nearly obstructing oxidative phosphorylation. Finally, we observed a gradual shift toward glycolysis in the presence of doxorubicin, etoposide, and galactosamine (Figure 4A). Nonetheless, it is noteworthy that doxorubicin treatment compromised the overall respiratory capacity. These findings underscore the intricate and diverse effects of senescence inducers on cellular bioenergetics.

**Figure 4.**
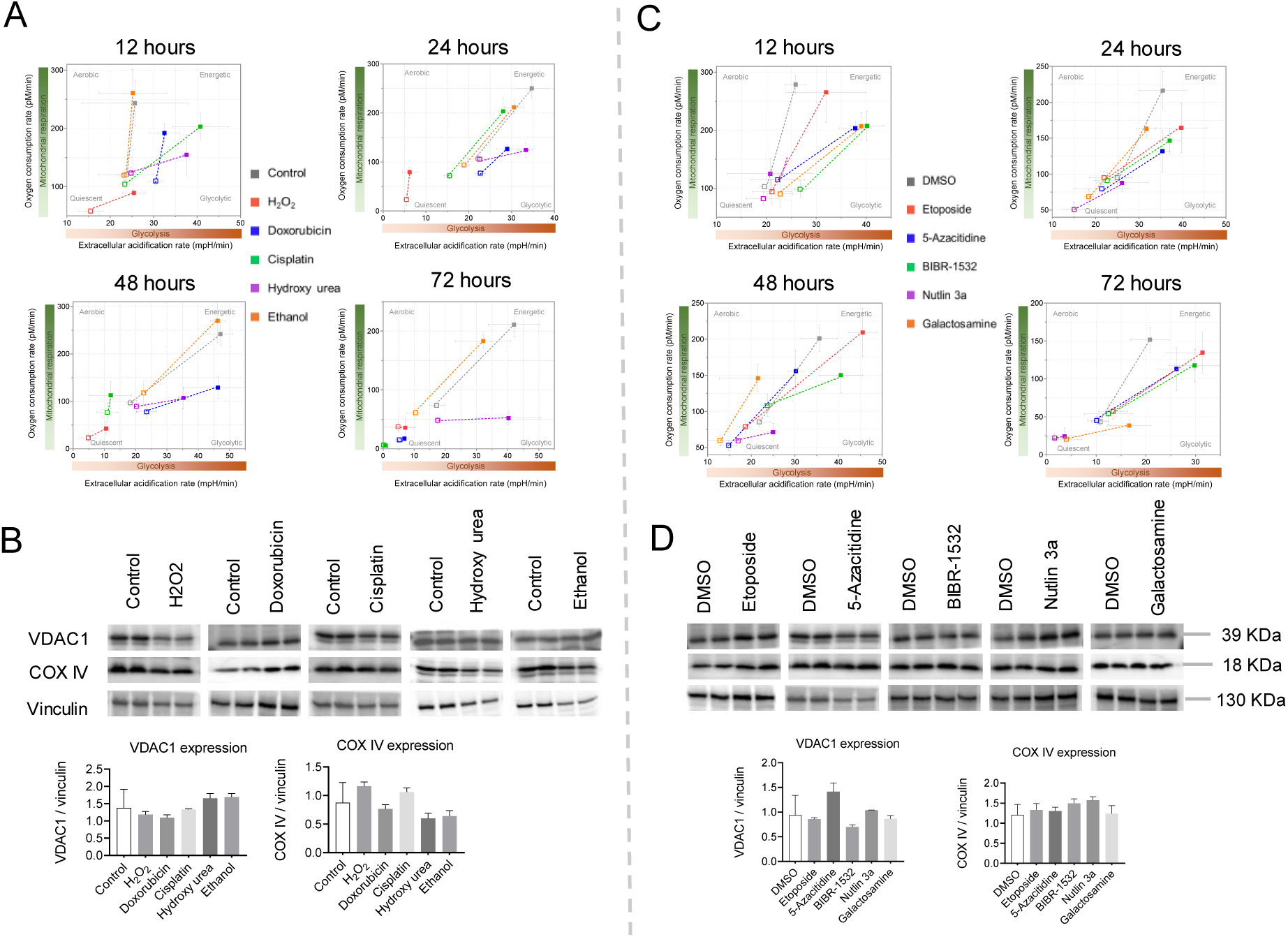
Senescence alters cellular bioenergetics without affecting the mitochondrial mass in primary mouse hepatocytes. A) Oxidative phosphorylation and glycolytic capacities measured by Seahorse flux analyzer. Basal and stressed levels were measured with 10 mM glucose, 2 mM pyruvate, and 1 mM glutamine as substrates. Error is shown as standard deviation. B) Representative Immunoblotting images of the proteins for assessing mitochondrial mass (VDAC1 and COX IV). Vinculin served as a loading control. The band intensity was quantified using Image J. Error is shown as standard deviation (n=3).

To discern the potential role of mitochondrial mass in the observed fluctuations of cellular bioenergetics, we conducted an evaluation of the expression levels of key mitochondrial proteins, namely VDAC1 and COX IV. Although slight variations were noted in their expression patterns, no significant changes were observed in mitochondrial mass across the treatments with senescence inducers (Figure 4B). This suggests that the observed changes in cellular bioenergetics were not primarily driven by changes in mitochondrial mass.

### 2.5 Key effects of senescence inducers on 2D hepatocyte culture are confirmed in hepatocyte-derived organoids

To ascertain that the limited proliferation of hepatocytes in a 2D culture does not compromise the senescence phenotype, we conducted additional experiments employing hepatocyte-derived organoids to validate our initial findings. Notably, we observed a more pronounced response to senescence inducers in these organoids compared to cells cultured in 2D, as evidenced by a heightened presence of senescence markers. Specifically, the application of senescence inducers promoted the phosphorylation of ɣH2A.X, a well-established marker associated with DNA damage, followed by the activation of p53 and p21. To evaluate the nuclear membrane’s integrity, we quantified the expression of lamin B1. Interestingly, all the senescence inducers resulted in a reduction of lamin B1 expression within the organoids relative to the control group (Figure 5A) (Figure S5). Moreover, the senescence inducers augmented the expression of various candidates associated with the senescence-associated secretory phenotype (SASP) within the organoids, surpassing the levels observed in 2D hepatocytes. Surprisingly, despite inducing senescence, all the senescence inducers caused a decrease in the expression of TGFβ, a pro-senescence cytokine (Figure 5B).

**Figure 5.**
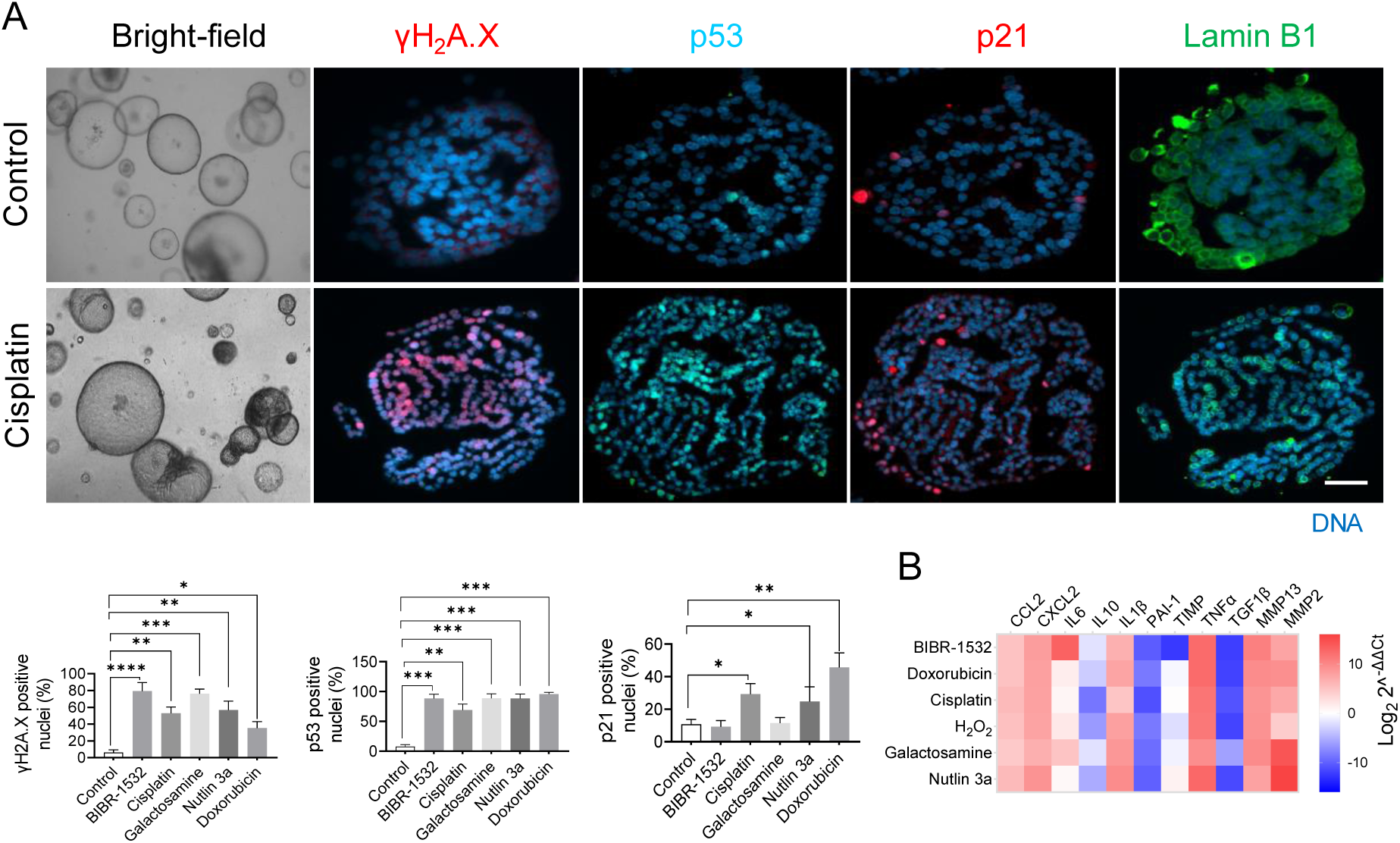
Effect of senescence inducers on hepatocyte-derived organoids. A) Represented images of senescence markers detected by immunofluorescence in hepatocytes-derived organoids treated with senescence inducers for 48 hours. The percentage of positive nuclei was plotted and shown in the bar graphs. Data were collected from three biological replicates (n=3) and two technical replicates. The mean of technical replicates was used for statistical analysis. Student’s t-test was used to compare the effect of each treatment with the respective control. Error is shown as standard deviation. (*P < 0.05, **P < 0.005, ***P < 0.0001, and ****P < 0.00001). B) Heat map representing the gene level expression of cytokines, chemokines, and matrix metalloproteases selected among the SASP candidates. GAPDH served as a reference gene and the data show the fold change over the control group. The data represent the mean of three biological (n=3) and two technical replicates.

These findings highlight the disparity in the extent of senescence markers observed between 2D cultures and hepatocyte-derived organoids. Nevertheless, the replication of the observed results from the 2D culture within the hepatocyte-derived organoids further confirms the validity and relevance of our initial findings.

### 2.6 Correlating senescence induction mechanisms with molecular pattern alterations

In order to summarize the findings and estimate the “net effect” of different stimuli on the strength of inducing cellular senescence in primary hepatocytes, we developed a scoring system based on 32 markers for senescence and apoptosis. Scoring considered ‘expected’ and ‘actual’ changes in markers

Score (+1): Expected increase, actual increase

Score (0): Expected increase/decrease, actual no change

Score (-1): Expected increase, actual decrease

The detailed table is provided in supplementary (Table S1). After scoring, the arithmetic summation was calculated and plotted as a bar graph (Figure 6A&B). Doxorubicin was the most potent senescence inducer, followed by cisplatin and H2O2, not affecting apoptosis. Unexpectedly, telomerase inhibition scored zero, with half of the markers aligning as expected and half opposing. Nevertheless, telomerase inhibition activated apoptosis *via* Bax/Bcl2.

**Figure 6.**
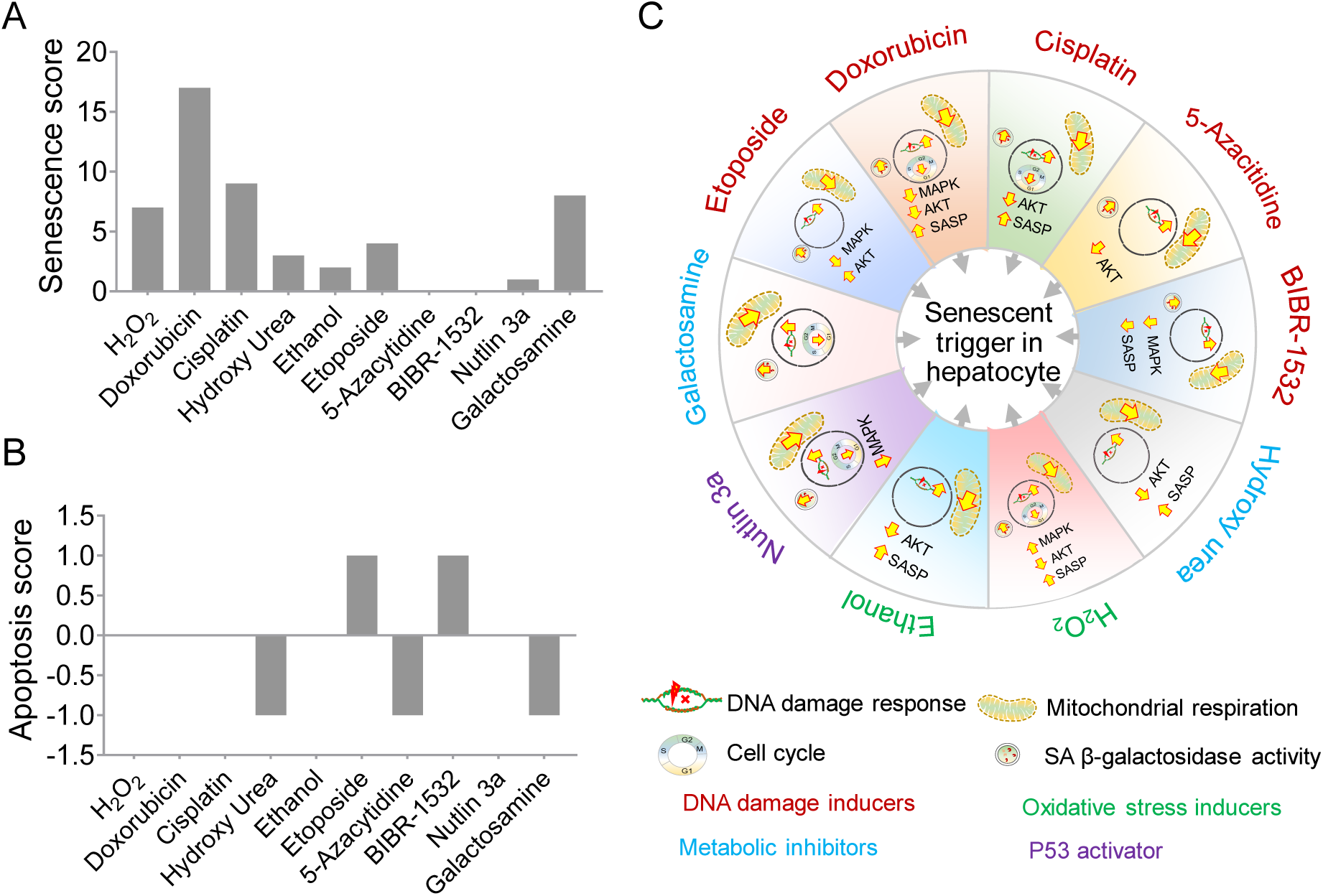
Senescence and apoptosis scoring of senescence inducers. A) The senescence score was calculated based on the relative change in the parameter in a drug-treated group compared to the untreated group. B) Apoptosis score was calculated based on the relative change in the parameter in a drug-treated group compared to the untreated group. C) Summary of the major molecular changes and the mode of senescence induction in PMH.

Next, we summarized senescence marker changes and inducer mechanisms. Markers included DNA damage response, cell cycle, mitochondrial dysfunction, SA-β-gal, SASP, and kinases. All inducers induced DNA damage response and mitochondrial dysfunction, independent of MAPK/AKT. Notably, only telomerase inhibition reduced SASP candidate expression. Our study revealed inconsistencies in the MAPK-SASP link: Doxorubicin reduced MAPK but raised SASP, while H_2_O_2_ increased both (Figure 6C, Figure S6).

## 3. Discussion

In the present study, we explored the molecular heterogeneity of senescence and revealed specific changes in molecular patterns and cellular bioenergetics in PMH. We report that each senescence inducer triggers a unique combination of senescence markers and this diversity is independent of the class of senescence inducer. Doxorubicin was the most potent senescence inducer, followed by cisplatin and H_2_O_2_. All inducers activated DNA damage response and mitochondrial dysfunction, independent of MAPK/AKT.

The liver consists of different cell types and each may respond differently to the senescence stimulus. Senescence in non-parenchymal cells (e.g. HSC) may restrict their activation (which should be investigated in further studies) and hence limit the liver fibrosis whereas senescent parenchymal cells (hepatocytes) produce pro-inflammatory and pro-fibrotic factors that activate HSCs, leading to the deposition of extracellular matrix proteins and the development of fibrosis ^23 24 25 26^. However, it is crucial to investigate senescence in a cell type-specific manner, as the consequences can vary significantly. For instance, while senescent hepatic stellate cells have been shown to suppress liver fibrosis, senescent hepatocytes may actually promote fibrosis within the liver. Moreover, the number of senescent hepatocytes resulting from telomere shortening has been found to exhibit an inverse correlation with the progression of NAFLD ^27^. In a separate study, Tabibian et al. (2014) discovered that senescence induced by bile acids in cholangiocytes leads to primary sclerosing cholangitis through the activation of N-Ras ^28^. Although the precise cause of immunosenescence remains elusive, the presence of senescent peripheral lymphocytes has been observed in patients with chronic viral hepatitis ^29^ as well as cirrhosis ^30^.

Although *in vitro* models have limitations in terms of lack of multicellular complexity and low physiological relevance, they have advantages in terms of accessibility and reproducibility. Since our study focuses on specifically one cell type, the *in vitro* cell culture serves as an ideal model to identify the senescent potential of different stress inducers. We have also used a hepatocyte-derived organoids system that helped us rule out the limited proliferation of PMH in 2D culture.

Since its discovery, and in general, senescence is considered to be caused by chronic stress, however, there are significant reports finding the activation of the senescence process acutely, usually 12-48 hours of stress ^31 32^. Our study also strengthens the reports of senescence induction by acute stressors and discovers that the dosage and time of exposure of the stressors are the keys to driving the cells toward senescence.

Despite the well-characterized hallmarks of the senescence phenotype ^33^, at the molecular level, no two senescence inducers had shown the same pattern of changes in all the measured senescence markers. Increased activity of Senescence Associated β-Galactosidase (SA-β-Gal) is the most commonly used marker to identify senescence and we found that all the senescence inducers used in the study increased its activity in PMH. However, PMH showed low specificity for SA-β-Gal as untreated cells also exhibited high activity that further elevated in a time-dependent manner. Therefore, subtle differences in senescence levels may not be accurately detected by this marker, specifically in PMH in 2D culture.

The DNA damage response and p53 pathway activation are key events in senescence. Doxorubicin emerged as a potent inducer, promoting DNA damage by phosphorylation of H2AX and activating the p53 pathway, leading to p21 upregulation, and promoting cell cycle arrest. Etoposide, despite not inducing H2AX phosphorylation, activated the p53 pathway, suggesting alternative mechanisms of DNA damage. H_2_O_2_ induced p53 activation, leading to p21 activation without significant H2AX phosphorylation, highlighting its distinct mode of action. These findings are particularly relevant to liver diseases characterized by DNA damage accumulation, such as chronic viral hepatitis and non-alcoholic fatty liver disease (NAFLD).

Cell cycle arrest is a prominent feature of senescence. Our results demonstrated that senescence inducers exerted diverse effects on cell cycle progression and proliferation markers. H_2_O_2_ induced phosphorylation of cdc2 and increased cyclin A2 levels, indicating attempts at proliferation despite obstacles in the cell cycle and DNA replication. In contrast, doxorubicin and cisplatin caused robust cell cycle arrest by downregulating multiple markers. Hydroxyurea and ethanol displayed imbalances in cell cycle regulation and DNA replication. Etoposide and 5-azacitidine affected specific cell cycle markers, potentially influencing cell growth. BIBR-1532, Nutlin 3a, and galactosamine showed distinct effects on cell cycle and proliferation markers, suggesting disruption of cell cycle progression.

The SASP is a crucial component of senescent cells, contributing to tissue microenvironment remodeling. Our findings revealed unique SASP profiles induced by different senescence inducers. Despite doxorubicin and etoposide both targeting topoisomerase II, they exhibited distinct SASP profiles. Cisplatin and 5-azacitidine, affecting DNA integrity, elicited specific SASP profiles. H_2_O_2_ and ethanol, inducing oxidative stress, showed differential SASP expression patterns. Galactosamine and hydroxyurea, interfering with nucleotide pools, influenced SASP expression. The p53 activation with Nutlin 3a affected SASP expression, highlighting the regulatory role of p53 in SASP modulation.

A cell undergoes a huge bioenergetic adaptation during senescence. Contradictory reports exist regarding whether cells rely more on glycolysis, oxidative phosphorylation, or both. Studies have shown that senescent lymphoma cells induced by DNA damage (cyclophosphamide) exhibited increased energy production through elevated glycolysis, lactate production, and oxidative phosphorylation in mice ^34^. Similarly, replicative senescent human diploid fibroblasts had elevated glucose consumption and lactate production ^35 36^. Recent metabolomic approaches on extracellular metabolites released by senescent fibroblasts supported a shift toward glycolysis compared to young cells ^19^ although this shift was not consistently observed in different cell types as observed by Delfarah et al., who found unaltered glucose consumption and lactate secretion in senescent human mammary epithelial cells ^37^. Conversely, numerous studies reported that a high glycolysis rate, accompanied by elevated OXPHOS activity, was specific to oncogene-induced senescence ^22 34 38^. Thus, it is evident that changes in cellular bioenergetics during senescence vary depending on the cell type and the underlying cause of senescence. Our study further supports this notion, as we observed significant variability in the pattern of cellular bioenergetics among senescent hepatocytes in mice exposed to different senescence-inducing factors. In general, DNA damage-induced senescence initially increased oxidative phosphorylation (within 24 hours of treatment) but shifted towards glycolysis at later time points, even while other senescent markers were present throughout. This shift may be attributed to the energy-demanding DNA repair mechanism triggered by DNA damage, which is complemented by increased oxidative phosphorylation. However, once the cell surpasses the recovery threshold, glycolysis becomes the primary energy source. This pattern was not observed in other senescence inducers where DNA damage was not the primary cause of senescence induction.

In conclusion, our study reveals the diverse effects of senescence inducers on molecular patterns and cellular bioenergetics in primary mouse hepatocytes (PMH). Each inducer triggers a unique combination of senescence markers, independent of their class. Senescence plays a role in various liver pathologies, but its consequences vary depending on the cell type involved. Our findings challenge the notion that senescence is solely caused by chronic stress, as acute stressors can also induce senescence. We observed distinct molecular changes, DNA damage response, and cell cycle alterations among senescence inducers. The senescence-associated secretory phenotype (SASP) showed unique profiles induced by different inducers, impacting tissue microenvironment remodeling. Cellular bioenergetics exhibited significant variability, with a shift from oxidative phosphorylation to glycolysis observed in DNA damage-induced senescence. These findings enhance our understanding of senescence induction, molecular heterogeneity, and interplay among their patterns.

## 4. Methods

### 4.1 Isolation of primary hepatocytes

The isolation of primary hepatocytes was carried out using a well-established method involving a series of steps. Firstly, the mouse liver (specifically, C57B6/J mice aged 18-23 weeks) was subjected to Ca^2+^-free perfusion to remove the blood and non-parenchymal cells from the liver tissue, followed by collagenase perfusion.

Hepatocytes were further purified (obtained >90 % viability) by percoll gradient as described ^39^. Subsequently, the isolated and purified hepatocytes were seeded onto cell culture plates or glass dishes coated with collagen type 1 (5 µg/cm2) at a density appropriate for the experimental requirements. The culture medium used for sustaining the hepatocytes was William’s E-medium supplemented with 2% fetal bovine serum (FBS), 2 mM L-glutamine, 5% Penstrep (penicillin-streptomycin), and hepatocyte growth supplements (CM4000) obtained from Thermo Fisher Scientific. This complete medium provided the necessary nutrients, growth factors, and antibiotics to support the growth and maintenance of the primary hepatocytes *in vitro*.

### 4.2 Senescence associated β-Galactosidase activity measurement

To investigate the effects of senescence inducers on hepatocytes, cells were treated with the respective senescence inducers for varying durations, including 12 hours, 24 hours, 48 hours, and 72 hours. Following treatment, the cells were incubated in a controlled environment of a humidified incubator set at 37°C and 5% CO_2_, providing optimal conditions for cellular growth and activity. Hepatocytes were treated with senescence inducers for respective times (12 h, 24 h, 48 h, and 72 h) and incubated in a humidified incubator at 37°C and 5% CO_2_. Senescence-associated β-Galactosidase (SA β-gal) activity was measured in the hepatocytes using Senescence β-Galactosidase Staining Kit (Cell Signaling Technology-9860, Danvers, MA, USA) as per manufacturer’s instructions. The signals were imaged with an inverted microscope Axio Observer 7 (Carl Zeiss Microscopy, White Plains, NY, USA). The signals were quantified with ImageJ software ^40^.

### 4.3 Immunofluorescence

Hepatocytes were cultured in glass bottom dishes (Cellvis-D35-14-0-N, Mountain View, CA, USA) and treated with senescence inducers for 24 hours. Cells were fixed with 4% paraformaldehyde for 15 minutes followed by permeabilization with 0.2% Triton X-100 for 15 minutes at room temperature. Antigens were blocked with 2% normal goat serum and 1% bovine serum albumin (BSA) for 1 hour in a humidified chamber at room temperature to prevent non-specific binding of the antibodies. Then, cells were incubated with primary antibodies (table S2) diluted in Antibody Diluent (Thermo Fisher Scientific-003218, Waltham, MA, USA) overnight (16 h) in a humidified chamber at 4 °C. Next day, the primary antibodies were washed off and the samples were incubated with fluorescently labeled secondary antibodies (table S2). Then the DNA was stained with DAPI (Sigma-Aldrich-D9542, St. Louis, MO, United States) and the F-actin was stained with Phalloidin-iFluor 488 (Abcam-ab176753, Cambridge, United Kingdom). Samples were mounted with VectaMount^®^ AQ Aqueous Mounting Medium (Vector laboratories-H-5501-60, Newark, CA, USA) and imaged with Axio Observer 7 (Carl Zeiss Microscope). Fluorescence intensity or the number of nuclei was measured with Image J software.

### 4.4 Tissue Lysis and Immunoblot Analysis

Hepatocytes were homogenized in RIPA buffer (150 mM NaCl, 1% NP-40, 0.5% Na-deoxycholate, 0.1% SDS, and 50 mM Tris-HCl pH 7.4). To ensure the preservation of protein integrity and activity, the RIPA buffer was supplemented with protease and phosphatase inhibitors (Thermo Fisher Scientific-78444). Protein concentration was measured with the Pierce^TM^ BCA assay (Thermo Fisher Scientific-23227). Equal amounts of proteins were separated by sodium dodecyl sulfate-polyacrylamide gel electrophoresis (SDS-PAGE) and transferred to a nitrocellulose membrane, blocked for 1 h with 5% non-fat milk or BSA, then incubated overnight at 4 °C with primary antibodies (table S2). After incubation with peroxidase-conjugated secondary antibody (Thermo Fisher Scientific, Rockford, IL, USA), signals were revealed with enhanced chemiluminescence (ChemiDoc MP, BioRad, Hercules, CA, USA). The signal intensity was quantified with Image J software.

### 4.5 Isolation of Total RNA and Quantitative PCR

Total RNA was extracted from the samples using RNA-Solv^®^ Reagent following the manufacturer’s instructions (Omega Bio-Tek-R6830-02, Norcross, GA, USA) and subsequently stored at -80 °C to ensure its stability. For reverse transcription, one microgram of the extracted RNA was converted into complementary DNA (cDNA) using Reverse Transcriptase (Roche-4897030001, Basel, Switzerland), a high-quality enzyme known for its reliable performance. To facilitate the amplification of specific target genes, gene primers were manually designed and synthesized by Eurofins Scientific (Luxembourg). The primer details, including the sequences and target genes, can be found in the accompanying table (table S3). Amplification of the cDNA samples was carried out using PowerUp^TM^ SYBRTM Green Mastermix (Thermo Fisher Scientific-A25742, Rockford, IL, USA). The real-time PCR reactions were performed using the QuantStudio™ 3 Real-Time PCR System (Thermo Fisher Scientific).

To analyze the gene expression data, the ΔCt (delta Ct) method was employed, whereby the Ct values of the target genes were normalized to the expression levels of the housekeeping gene, Glyceraldehyde 3-phosphate dehydrogenase (GAPDH). This normalization approach allows for the comparison and interpretation of relative gene expression levels across different samples. The resulting values were reported as fold changes, indicating the increase or decrease in expression relative to the control samples. The fold changes were calculated using the formula 2^(-ΔΔCt), which provides a quantitative measure of the relative changes in gene expression levels compared to the control group.

### 4.6 Measurement of the rate of glycolysis and oxygen consumption

The assessment of oxygen consumption rates (OCR) and extracellular acidification rate (ECAR) was conducted using a Seahorse Extracellular Flux (XF)-96 Bioanalyzer from Agilent Technologies. Primary hepatocytes and hepatocyte-derived organoids were utilized for this analysis. To initiate the experiment, hepatocytes were seeded at a density of 10,000 cells per well in 96-well plates that were pre-coated with collagen type I at a concentration of 5 µg/cm^2^. Following cell seeding, the hepatocytes were treated with specific senescence inducers for varying durations of time (12 hours, 24 hours, 48 hours, and 72 hours). Throughout the experimental period, the cells were incubated in a humidified incubator at a temperature of 37°C and a CO_2_ concentration of 5%.

For the OCR and ECAR measurements, the culture medium was replaced with XF basal medium from Agilent Technologies, supplemented with 10 mM glucose, 1 mM pyruvate, and 1 mM glutamine. Subsequently, the cells were incubated for 1 hour at 37°C in a non-humidified incubator without CO2. In order to analyze the OCR and ECAR, specific inhibitors including oligomycin (2 µM), FCCP (1 µM), rotenone (1 µM), and antimycin (1 µM) were titrated and delivered through injection at designated ports A, B, and C of the Bioanalyzer. The resulting data were then normalized to account for the total number of cells present in each well.

In the case of hepatocyte-derived organoids, pre-developed organoids were seeded in 20 µl of matrigel within the wells of a 96-well plate and allowed to grow for a period of 3 days. Subsequently, the organoids were treated with the respective senescence inducers for different durations (48 hours and 96 hours) and maintained in a humidified incubator at a temperature of 37°C and a CO2 concentration of 5%. Following the incubation period, the culture medium was replaced with XF basal medium supplemented with 10 mM glucose, 1 mM pyruvate, and 1 mM glutamine. Similar to the hepatocytes, the organoids were then incubated for 1 hour at 37°C in a non-humidified incubator without CO2. The inhibitors oligomycin (6 µM), FCCP (4 µM), rotenone (3 µM), and antimycin (3 µM) were titrated and injected at ports A, B, and C of the Bioanalyzer to assess OCR and ECAR.

### 4.7 Culture of HepOrgs

As described previously ^41^, 50 µl matrigel containing 5000 hepatocytes was plated in the center of 12-well glass-bottom chambers or 24-well plates. After a 5-min incubation for matrigel polymerization, 300 µl or 500 µl of complete medium (DMEM-F12) containing the growth factors N-acetylcysteine (1.25mM), mEGF (50ng/ml), R-spondin 1 (15%), gastrin (10 nM), hFGF10 (100 ng/ml), hHGF (25 ng/ml), nicotinamide (10nM), FGF7 (50ng/ml), Rho inhibitor (10 uM), CHIR99021 (3µM) and TGFBi (1µM) supplemented with B27 (without vitamin A) was added. The culture medium was replaced every 3 days until complete organoids had developed for 15 days.

On day 15, the medium was aspirated, 500 µl AdvDMEM+++ was added to each well, and the hepatocyte organoids were mechanically broken using a P1000 tip. Organoids were pipetted up and down 20 times with a glass Pasteur pipette (230 mm) and vigorously 20 times with a 1 ml pipette. After centrifugation, a sufficient amount of matrigel was added and the organoids were seeded at a 1:4 ratio.

### 4.8 Exposure of HepOrgs to senescence inducers and IF staining

Organoids from each well were exposed to different senescence inducers with pre-optimized dosages of senescence inducers for 48 h. After 48 hours, HepOrgs were subsequently washed twice with PBS, fixed with 4% paraformaldehyde for 20 minutes, and embedded in paraffine. 4.5-µm sections were deparaffinised, washed with PBS/glycine solution, and permeabilized with 0.5% Triton X 100 in PBS for 10 minutes on a shaker. Plates were then carefully washed with immunofluorescence wash buffer (IF) and blocked with 10% goat serum in the IF wash buffer for 2 hours on a shaker. Primary antibodies (table 3) were applied overnight at 4 °C. After washing 3 times with IF wash buffer for 10 minutes on a shaker, the secondary antibodies were applied 1:500 for 1 hour at room temperature. Slides were washed and incubated with DAPI (1:10,000) in 1× PBS for 10 minutes. Slides were mounted with Histokitt mounting medium (Carl Roth, Germany). Images of the HepOrgs were acquired using a Zeiss observer at 20X magnification.

### 4.9 Statistical Analysis

Data are presented as the mean values ± standard deviations (SD) or standard error mean (SEM). Statistical comparisons were made between control and senescence inducer-treated hepatocytes. The statistical test used for each dataset analysis is mentioned in the figure’s legend. A p-value of ≤ 0.05 was considered statistically significant.

## Supporting information

Kumar et al 2023 supplementary information

## Author Contributions

Conceptualization (PK, CE), Methodology (PK, MH), Investigation and data curation (PK, MH), Writing – original draft (PK, CE), Writing – review & editing (PK, MH, CE, FT), Formal Analysis (PK, MH), and Supervision (CE, FT) Funding acquisition (CE). All authors have read and agreed to the published version of the manuscript.

## Data Availability Statement

Specific data and materials described in this study will be made available upon request.

## Acknowledgments

We are thankful for technical support from the Department of Hepatology and Gastroenterology, Charité - Universitätsmedizin Berlin (CVK). We also thank Prof. Dr. med. Clemens A. Schmitt, Dr. Animesh Bhattacharya, Charité - Universitätsmedizin Berlin (CVK), and Max-Delbrück-Center for Molecular Medicine for providing the support with the Seahorse instrument.

