## Supplementary material for "Delineating the heterogeneity of senescence-induced-functional alterations in hepatocytes": Kumar et al 2023 supplementary information: Kumar et al 2023_supplementary information.pdf

**S1. Optimization of dosage of senescence inducers by SA-β-Gal activity assay in primary mouse hepatocytes**

To assess the effects of various senescence inducers on SA-β-Gal activity in primary mouse hepatocytes (PMH), we first optimized the diseases and time of treatment. We categorized the inducers based on their mode of action and observed their effects on SA-β-Gal activity.

*Oxidative Stress Inducer:* Hydrogen peroxide treatment at concentrations of 50 μM and 200 μM for 12 to 72 hours resulted in a significant increase in SA-β-Gal activity, indicating the induction of senescence. However, treatment with a higher concentration of 500 μM exhibited cytotoxic effects, leading to the absence of detectable SA-β-Gal activity.

*DNA-Damaging Agents:* Doxorubicin treatment at concentrations of 0.1 μM, 1 μM, and 5 μM for up to 48 hours increased SA-β-Gal activity in a time-dependent manner, suggesting the activation of senescence pathways. However, prolonged treatment for 72 hours reduced SA-β-Gal activity, possibly due to cytotoxicity. Etoposide, a topoisomerase II inhibitor, showed dose- and time-dependent increases in SA-β-Gal activity at concentrations of 1 μM, 10 μM, and 50 μM, suggesting the activation of senescence pathways associated with DNA damage. Cisplatin, a platinum-based chemotherapeutic agent, induced a time-dependent increase in SA-β-Gal activity at concentrations of 1 μg/ml and 5 μg/ml. Conversely, treatment with 20 μg/ml exhibited cytotoxic effects, resulting in lower SA-β-Gal activity.

*Metabolic Stress Inducers:* Hydroxyurea treatment at concentrations of 10 μM, 100 μM, and 500 μM increased SA-β-Gal activity in a time-dependent manner, indicating the induction of senescence through replication stress. Galactosamine treatment at a concentration of 0.1 mM showed a time-dependent increase in SA-β-Gal activity, indicating the induction of senescence. Interestingly, treatment with higher concentrations of 1 mM and 10 mM initially led to increased SA-β-Gal activity for up to 24 hours, followed by cytotoxic effects and a lower level of SA-β-Gal activity.

*Epigenetic Modulator:* 5-Azacitidine treatment at concentrations of 1 μM, 5 μM, and 20 μM resulted in a relatively lower increase in SA-β-Gal activity compared to other inducers. However, prolonged treatment for 72 hours with any concentration exhibited cytotoxic effects, suggesting a potential role in both senescence induction and cytotoxicity.

*Telomerase Inhibitor:* BIBR-1532 treatment at concentrations of 5 mM and 20 mM showed a time-dependent increase in SA-β-Gal activity, indicating the induction of senescence by telomerase inhibition. Notably, treatment with 100 mM exhibited cytotoxic effects and lower SA-β-Gal activity.

*p53 Activator:* Nutlin 3a treatment at concentrations of 1 μM and 10 μM induced a time-dependent increase in SA-β-Gal activity, suggesting the activation of senescence pathways mediated by p53. However, treatment with 50 mM showed cytotoxicity effects and a lower level of SA-β-Gal activity.

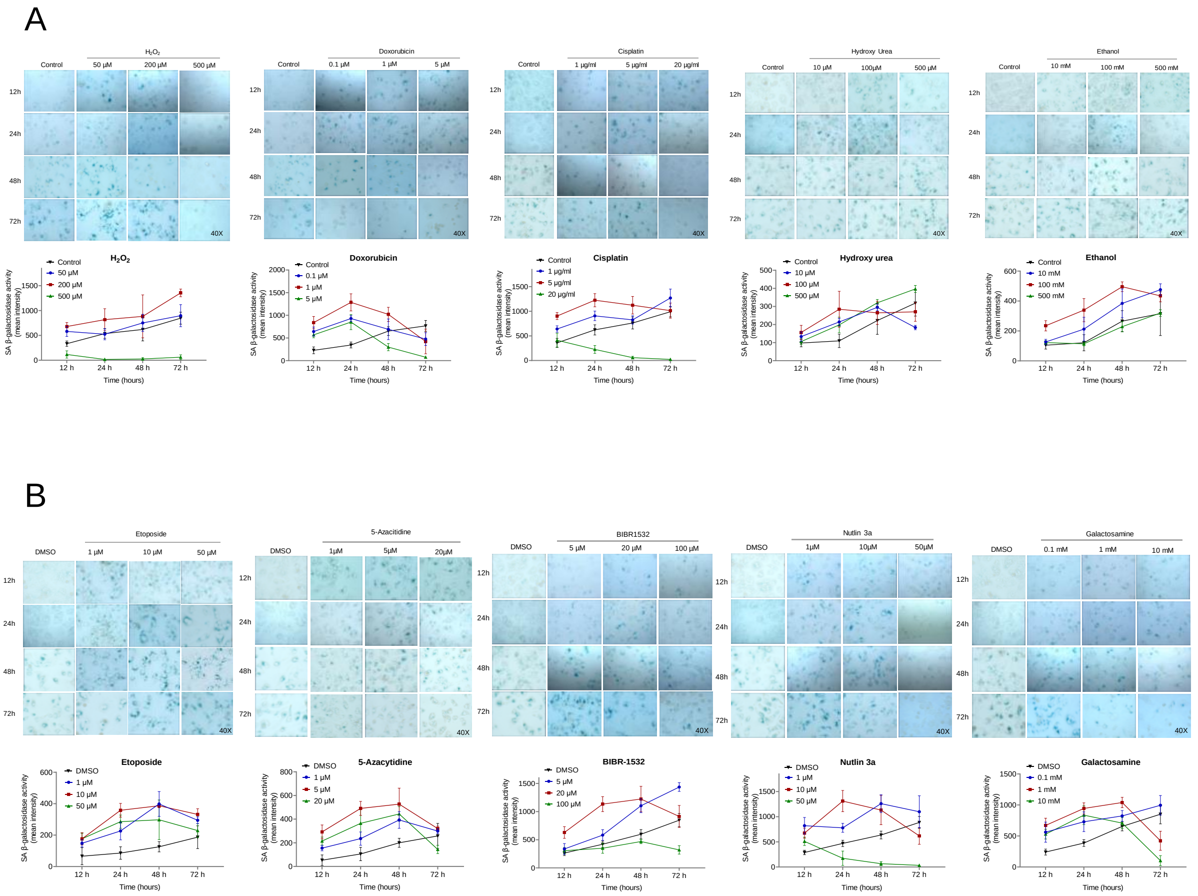


***Figure S1. Senescence inducers increased senescence-associated β-galactosidase activity in primary mouse hepatocytes***

*Treatment time (12h, 24h, 48h, and 72h) and dosage (three doses for each drug) of drugs were optimized and the representative images are shown. The intensity was quantified with Image J. Data represent three independent experiments. A) drugs dissolved in water and B) drugs dissolved in DMSO.*

### S2. Assessment of cell death markers under senescence inducers in primary mouse hepatocytes

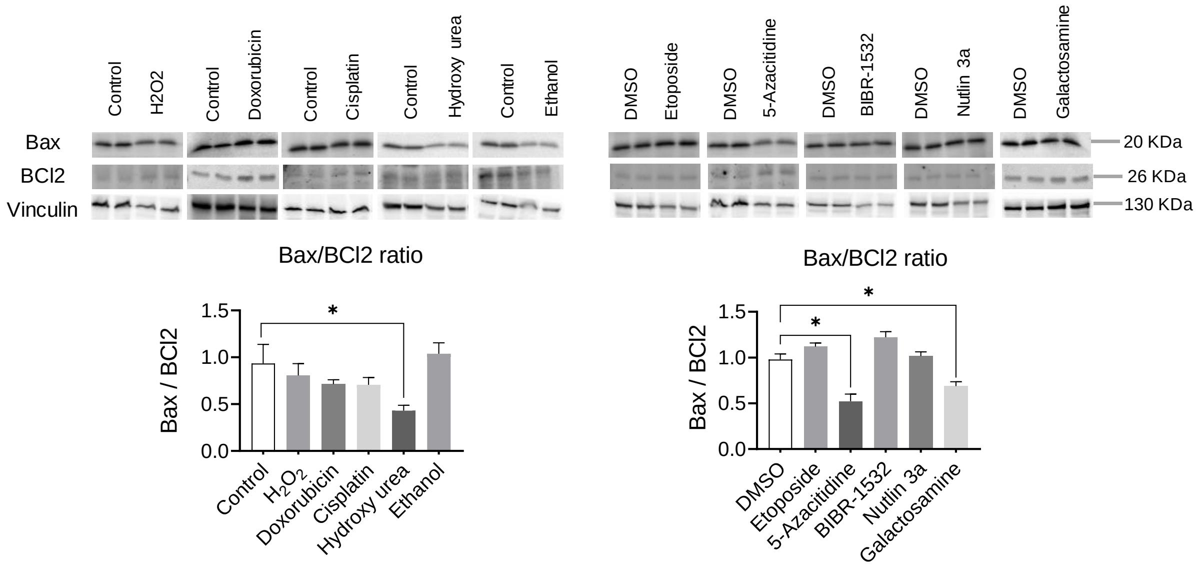


***Figure S2. Effect of senescence inducers on apoptotic markers in primary mouse hepatocytes***

*Representative Immunoblotting images of the effect of senescence inducers on cell cycle and proliferation markers in primary mouse hepatocytes treated for 24 hours in 2D culture. Vinculin served as a loading control. A) Drugs soluble in water and, B) relative quantification. C) Drugs soluble in DMSO and, D) relative quantification. Data is collected from four biological replicates (n=3). The band intensity was quantified using Image J. Student's t-test was used to compare the effect of treatment with the respective control. Error is shown as standard deviation. (*P < 0.05, **P < 0.001, and ***P < 0.001).*

**S3. Detailed images of senescence markers in primary mouse hepatocytes by immunofluorescence assessment.**

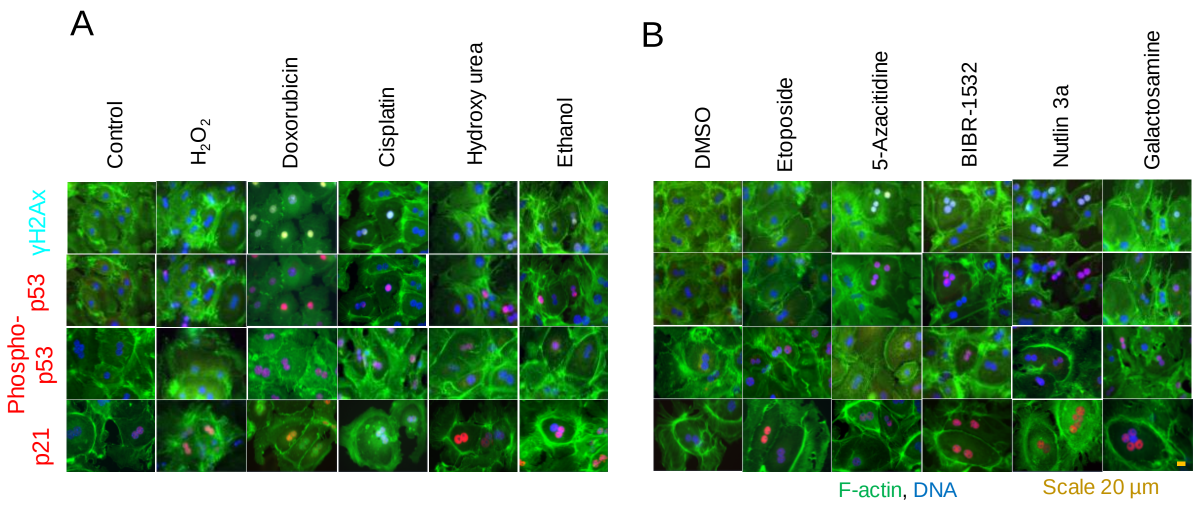


***Figure S3. DNA damage response is activated differently among senescence inducers:*** *Immunofluorescence images represent senescence-induced DNA damage response markers in primary mouse hepatocytes treated for 24 hours in 2D culture A) Drugs soluble in water and B) Drugs soluble in DMSO. Data is collected from four biological replicates (n=4) and five technical replicates.*

**S4. Effect of senescence inducers on nuclear membrane integrity in primary mouse hepatocytes.**

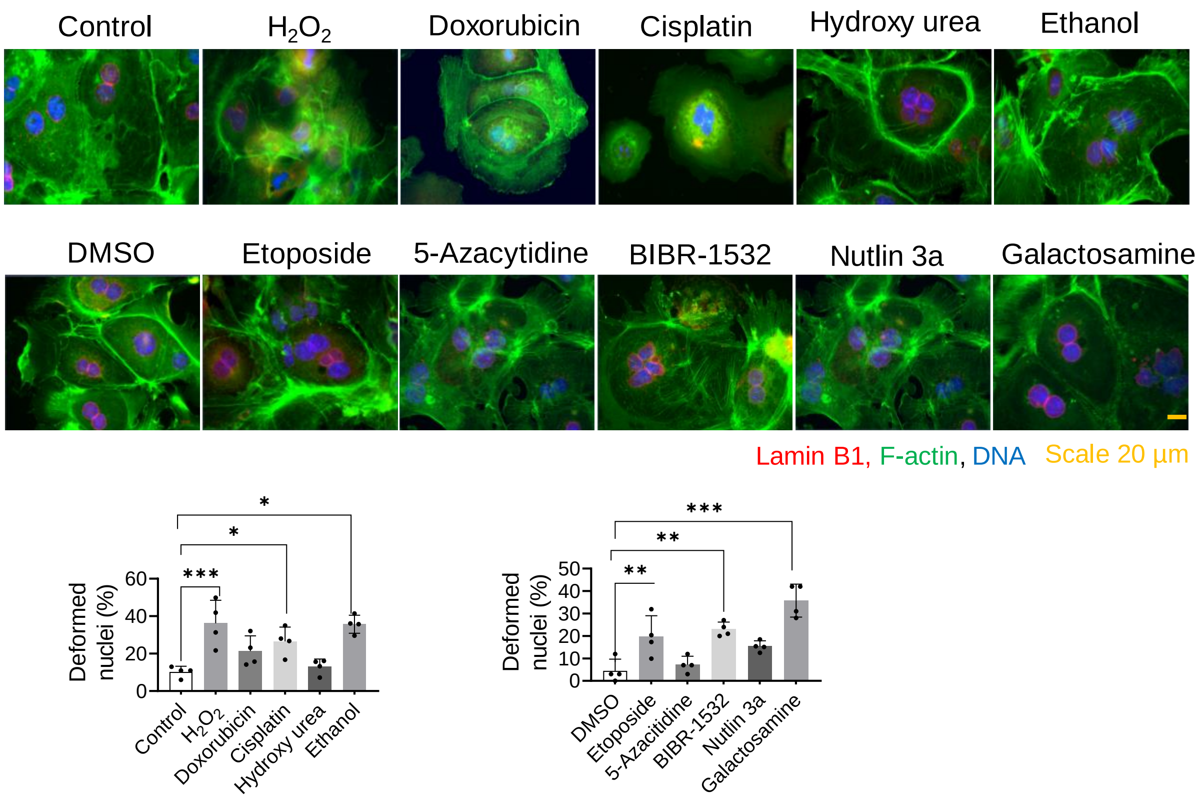


***Figure S4. Nuclear membrane integrity is lost in senescent primary mouse hepatocytes***

*Immunofluorescence images represent the nuclear membrane marker, lamin B1 in primary mouse hepatocytes treated for 24 hours in 2D culture. Bar graphs represent the percentage of deformed nuclei per field area. Data is collected from four biological replicates (n=4) and five technical replicates. The mean of technical replicates was used for statistical analysis. Student's t-test was used to compare the effect of treatment with the respective control. Error is shown as standard deviation. (*P < 0.05, **P < 0.001, ***P < 0.001, and ****P < 0.001).*

**S5. Detailed images of DNA damage response markers of senescence inducers treated hepatocyte-derived organoids.**

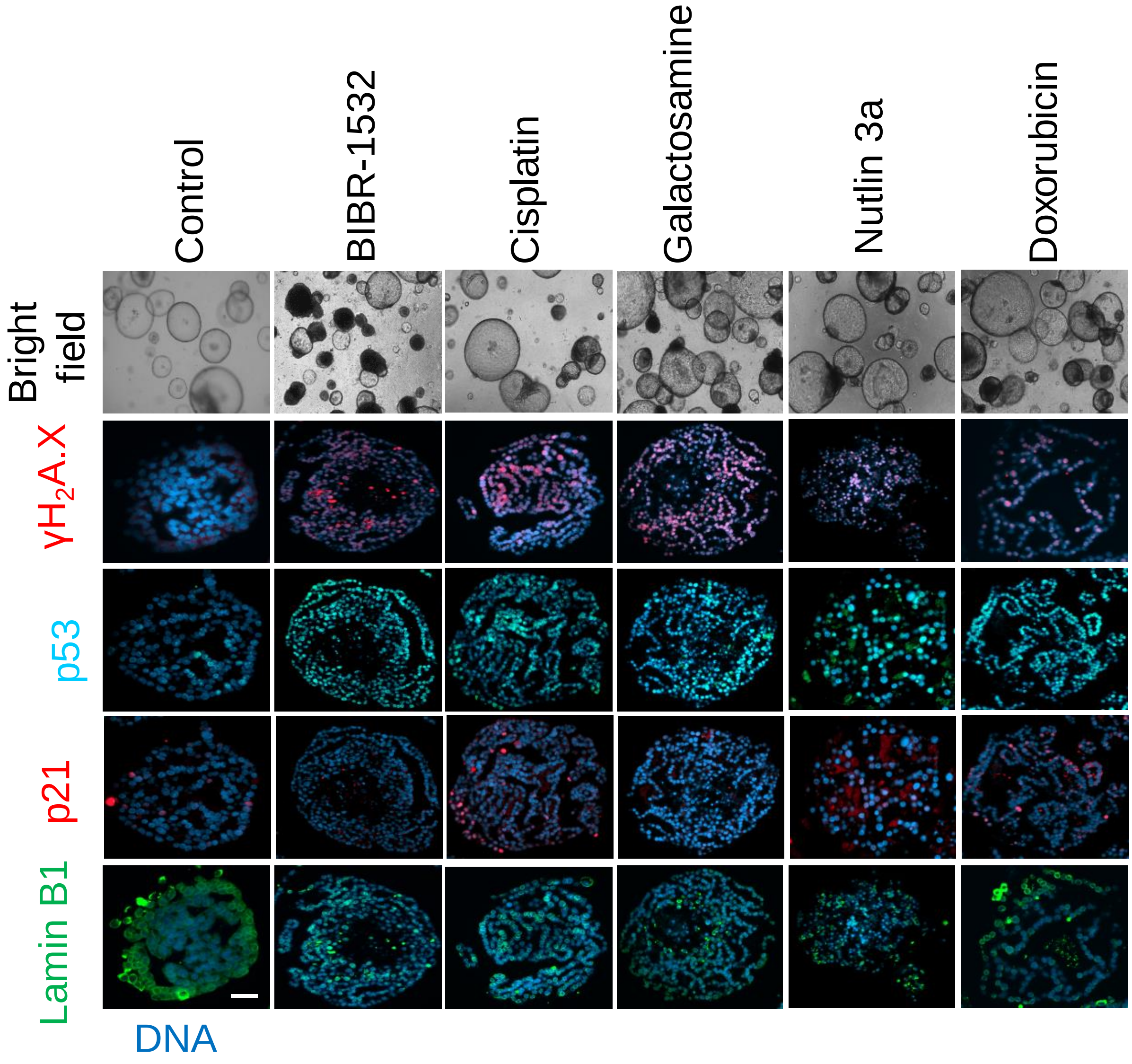


***Figure S5. DNA damage response is activated differently among senescence inducers:*** *Immunofluorescence images represent senescence-induced DNA damage response markers hepatocyte-derived organoids. The organoids were treated for 48 hours.*

### S6. Correlation among the senescence inducers and senescence markers

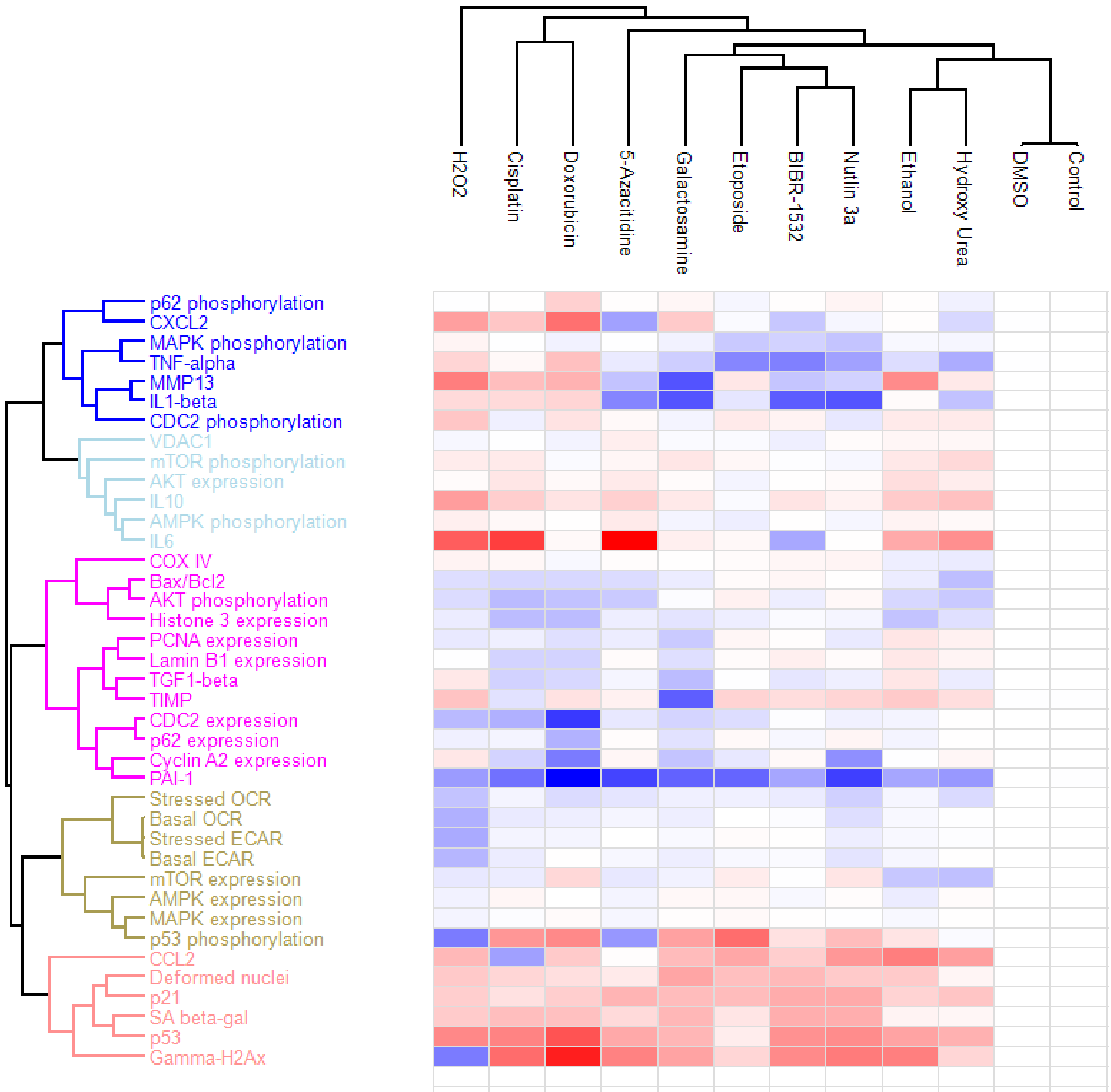


***Figure S6. Hierarchical clustering of the senescence inducers and their respective changes in the molecular markers.*** *Red color indicates increased and blue color indicates decreased changes in the markers compared to the respective control. The white color indicates no changes.*

**Table S1. Senescence and apoptosis score of senescence inducers**

| S.No. | Methods | Markers | $H_2O_2$ | Doxorubicin | Cisplatin | Hydroxy Urea | Ethanol | Etoposide | 5-Azacytidine | BIBR-1532 | Nutlin 3a | Galactosamine |
|---|---|---|---|---|---|---|---|---|---|---|---|---|
| 1 | SA β-galactosidase activity | | 1 | 1 | 1 | 0 | 0 | 0 | 1 | 1 | 1 | 1 |
| 2 | Immunoflorescence | γH2A.X | 0 | 1 | 1 | 0 | 1 | 0 | 1 | 1 | 1 | 1 |
| 3 | | p53 | 1 | 1 | 1 | 1 | 1 | 0 | 1 | 1 | 1 | 1 |
| 4 | | p_p53 | 0 | 1 | 1 | 1 | 1 | 1 | 0 | 0 | 1 | 1 |
| 5 | | p21 | 1 | 1 | 1 | 1 | 1 | 1 | 1 | 1 | 1 | 1 |
| 6 | | Deformed nuclei | 1 | 1 | 1 | 0 | 1 | 1 | 0 | 1 | 0 | 1 |
| 7 | Immunoblotting | Cyclin A2 expression | -1 | 1 | 1 | 0 | 0 | 1 | 0 | 0 | 1 | 1 |
| 8 | | CDC2 phosphorylation | -1 | 0 | 1 | 0 | 0 | -1 | 0 | 0 | 1 | 1 |
| 9 | | CDC2 expression | 1 | 1 | 1 | -1 | 0 | 1 | 1 | 0 | 0 | 1 |
| 10 | | Histone 3 expression | 1 | 1 | 1 | 1 | 1 | 0 | 0 | 0 | 0 | 1 |
| 11 | | Lamin B1 expression | 0 | 1 | 1 | 0 | -1 | 0 | 0 | 0 | 0 | 1 |
| 12 | | PCNA expression | 1 | 1 | 1 | 0 | -1 | 0 | 0 | 0 | 0 | 1 |
| 13 | qPCR (SASP) | CCL2 | 1 | 1 | -1 | 1 | 1 | 1 | 0 | 1 | 1 | 1 |
| 14 | | CXCL2 | 1 | 1 | 1 | -1 | 0 | 0 | -1 | -1 | 0 | 1 |
| 15 | | IL6 | 1 | 0 | 1 | 1 | 1 | -1 | 1 | -1 | -1 | 1 |
| 16 | | IL10 | 1 | 1 | 1 | 1 | 1 | -1 | 1 | 1 | 1 | 1 |
| 17 | | IL1β | 0 | 1 | 1 | -1 | -1 | -1 | -1 | 0 | 0 | 0 |
| 18 | | PAI-1 | -1 | 0 | -1 | -1 | -1 | -1 | 0 | -1 | 0 | -1 |
| 19 | | TIMP | 1 | 1 | -1 | 1 | 1 | 1 | 1 | 1 | 1 | -1 |
| 20 | | TNFα | 1 | 1 | 1 | -1 | -1 | -1 | -1 | 0 | -1 | -1 |
| 21 | | TGF1β | 1 | -1 | -1 | -1 | 1 | 0 | -1 | -1 | -1 | -1 |
| 22 | | MMP13 | 1 | 1 | 1 | 1 | 1 | 1 | -1 | -1 | -1 | 0 |
| 23 | Immunoblotting | AKT phosphorylation | -1 | -1 | -1 | -1 | -1 | 0 | -1 | -1 | 0 | 0 |
| 24 | | AKT expression | 0 | 0 | 1 | 1 | 1 | 0 | 1 | 0 | 0 | 0 |
| 25 | | MAPK phosphorylation | 0 | 0 | 0 | 0 | 0 | -1 | 0 | -1 | -1 | -1 |
| 26 | | MAPK expression | 0 | 1 | 0 | 0 | -1 | 1 | 0 | 0 | 0 | 0 |
| 27 | | mTOR phosphorylatior | 1 | 0 | 0 | 1 | 1 | 1 | 0 | 0 | -1 | 1 |
| 28 | | mTOR expression | -1 | 1 | -1 | -1 | -1 | 1 | -1 | 0 | 1 | 0 |
| 29 | Cellular bioenergetics | Basal ECAR | -1 | 0 | -1 | 0 | -1 | -1 | -1 | -1 | -1 | -1 |
| 30 | | Stressed ECAR | -1 | 1 | -1 | 1 | -1 | 1 | 1 | 1 | -1 | -1 |
| 31 | | Basal OCR | -1 | -1 | -1 | 0 | -1 | 0 | -1 | 0 | -1 | -1 |
| 32 | | Stressed OCR | -1 | -1 | -1 | -1 | -1 | -1 | -1 | -1 | -1 | -1 |
| | **Over all senescence score** | | **7** | **17** | **9** | **3** | **2** | **3** | **0** | **0** | **1** | **8** |
| | **Cell death score (Bax/BCl2 based)** | | **0** | **0** | **0** | **-1** | **0** | **1** | **-1** | **1** | **0** | **-1** |

**Table S2. Detailed information for antibodies used**

| Primary antibodies | | | | |
|---|---|---|---|---|
| **S.No.** | **Antibody** | **Company** | **Cat. Number** | **Dilution** |
| **1** | ɣH2A.X | Abcam | ab81299 | 1:400 for IF |
| **2** | p53 | Cell Signaling Technology | 2524 | 1:200 for IF, 1:1000 for WB |
| **3** | Phospho (S15) p53 | Cell Signaling Technology | 9284 | 1:200 for IF |
| **4** | p21 | Abcam | ab188224 | 1:400 for IF, 1:1000 for WB |
| **5** | Lamin B1 | Proteintech | 66095-1-Ig | 1:300 for IF, 1:5000 for WB |
| **6** | VDAC1 | Proteintech | 10866-1-AP | 1:100 for IF |
| **7** | Cyclin A2 | Abcam | ab181591 | 1:2000 for WB |
| **8** | Phospho CDC2 | Cell Signaling Technology | 9111 | 1:1000 for WB |
| **9** | CDC2 | Cell Signaling Technology | 28439 | 1:1000 for WB |
| **10** | Histone 3 | Cell Signaling Technology | 4499 | 1:2000 for WB |
| **11** | PCNA | Abcam | Ab29 | 1:1000 for WB |
| **12** | Vinculin | Merck millipore | CP74 | 1:1000 for WB |
| **13** | Phospho AKT | Cell Signaling Technology | 9271 | 1:1000 for WB |
| **14** | AKT | Cell Signaling Technology | 9272 | 1:1000 for WB |
| **15** | Phospho MAPK | Cell Signaling Technology | 4631 | 1:1000 for WB |
| **16** | MAPK | Cell Signaling Technology | 9212 | 1:1000 for WB |
| **17** | Phospho mTOR | Cell Signaling Technology | 5536 | 1:1000 for WB |
| **18** | mTOR | Cell Signaling Technology | 2972 | 1:1000 for WB |
| **19** | VDAC1 | Abcam | ab14734 | 1:1000 for WB |
| **20** | COX IV | Proteintech | 11242-1-AP | 1:5000 for WB |

| Secondary antibodies | | | | |
|---|---|---|---|---|
| **S.No.** | **Antibody** | **Company** | **Cat. Number** | **Dilution** |
| **1** | Goat anti-rabbit | BioRad | 1721019 | 1:10,000 for WB |
| **2** | Goat anti-mouse | BioRad | 1721011 | 1:10,000 for WB |
| **3** | Anti-rabbit AF-647 | Cell Signaling Technology | 4414 | 1:500 for IF |
| **4** | Anti-rabbit AF-555 | Cell Signaling Technology | 4413 | 1:500 for IF |
| **5** | Anti-mouse AF-647 | Cell Signaling Technology | 4410 | 1:500 for IF |
| **6** | Anti-mouse AF-555 | Cell Signaling Technology | 4409 | 1:500 for IF |

**Table S3. Primers for qPCR**

| | | Primers for qPCR | |
|---|---|---|---|
| **S.No.** | **Gene** | **Forward primer sequence (5’-3’)** | **Reverse primer sequence (5’-3’)** |
| **1** | CCL2 | GTGTTGGCTCAGCCAGATGC | GACACCTGCTGCTGGTGATCC |
| **2** | CXCL2 | CCCAGACAGAAGTCATAGCCAC | TGGTTCTTCCGTTGAGGGAC |
| **3** | IL6 | GCTACCAAACTGGATATAATCAGGA | CCAGGTAGCTATGGTACTCCAGAA |
| **4** | IL10 | GGCTGAGGCGCTGTCATCG | TCATTCATGGCCTTGTAGACACC |
| **5** | IL1β | GAGCTGAAAGCTCTCCACCTC | CTTTCCTTTGAGGCCCAAGGC |
| **6** | PAI-1 | GACACCCTCAGCATGTTCATC | AGGGTTGCACTAAACATGTCAG |
| **7** | TIMP | CCAGAACCGCAGTGAAGAGT | GTACGCCAGGGAACCAAGAA |
| **8** | TNFα | ACCACGCTCTTCTGTCTACTGA | TCCACTTGGTGGTTTGCTACG |
| **9** | TGF1β | ATTCAGCGCTCACTGCTCTT | ATGTCATGGATGGTGCCCAG |
| **10** | MMP13 | CCTTCTGGTCTTCTGGCACAC | GGCTGGGTCGTCACACTTCTCTGG |
| **11** | MMP2 | CAACGGTCGGGAATACAGCAG | CCAGGAAAGTGAAGGGGAAGA |
| **12** | HPRT1 | CAAACTTTGCTTTCCCTGGT | TCTGGCCTGTATCCAACACTTC |
| **13** | GAPDH | TGTTGAAGTCACAGGAGACAACCT | AACCTGCCAAGTATGATGACATCA |
